# Large-scale lensless microscopy with fast acquisition and region-wise focusing

**DOI:** 10.1101/2023.08.05.551428

**Authors:** You Zhou, Weizhi Song, Linyuan Wu, Lin Fan, Junjia Wang, Shaowei Jiang, Zhan Ma, Bo Xiong, Xun Cao

## Abstract

The imaging field of view (FOV) of lensless microscope is consistent with the size of image sensor in use, enabling the observation of sample areas larger than 20 mm^2^. Combined with high-performance and even super-resolution phase retrieval algorithms, micron and sub-micron resolution can be achieved, ultimately realizing wide-field and high-resolution imaging performance simultaneously. However, high-throughput lensless imaging poses significant challenges in terms of rapid data acquisition and large-scale phase retrieval. Additionally, when observing biological samples over a large FOV, the focus plane often exhibits inconsistency among different regions, necessitating further parameter calibration. In this study, we propose a fast acquisition and efficient reconstruction strategy for coherent lensless imaging based on a multi-height imaging model. Multiple measurements are manually modulated using an axial translation stage and continuously captured by an image sensor, facilitating rapid data acquisition within seconds and requiring no hardware synchronization. The efficiency and accuracy of phase retrieval are enhanced through precise parameter calibration algorithms, as well as techniques such as region-wise parallel computing and region-wise auto-focusing. Experimental results demonstrate 7.4×5.5 mm^2^ FOV and 1.55 μm half-pitch resolution imaging of human skin and lung tumor sections with region-wise focusing, requiring only an approximate 0.5-s acquisition time and 44-s reconstruction time. Furthermore, by incorporating the pixel super-resolution principle, the 1.10 μm half-pitch imaging resolution is demonstrated in full-FOV peripheral blood smears without additional data required, beneficial to the identification of hollow shape and segmentation of blood cells.

## 1 Introduction

High-throughput biological imaging has gained significant advancements in applications such as the observation of brain activity [1, 2], leucocyte trafficking [3], tumor metastasis [4], and large cell populations [5, 6]. Various high-throughput microscopic techniques have emerged, including lensless microscopy [7-9], array microscopy [2, 10-13], multiscale optical microscopy [1, 14-16], Fourier ptychographic microscopy (FPM) [7, 17, 18], and whole-slide scanning microscopy [19-21]. Among existing techniques, lensless imaging offers advantages such as low cost, system compactness, and portability, making it an important alternative for high-throughput optical microscopy and screening in point-of-care diagnostics [7, 8, 22-24]. In recent years, several advanced lensless imaging models and algorithms have been proposed, further enhancing the performance and expanding the applicability [9, 25-28]. Daloglu et al. propose a high-throughput and label-free imaging technique for three-dimensional (3D) lensless imaging of spermatozoon locomotion [25]. Adams et al. report the lensless FlatScope via an amplitude mask for 3D fluorescence imaging [26] and further employ an optimized phase mask to realize the lensless Bio-FlatScope for in vivo imaging of mouse brain and human oral mucosa [9]. Guo et al. design a depth-multiplexed ptychographic microscopy for parallel imaging of multiple stacked samples at high speed [27]. Jiang et al. propose a high-throughput ptychographic cytometry using a blood-coated image sensor for lensless acquisition and a modified Blu-ray drive for diversity modulation [28].

Lensless imaging eliminates the need for any optical lenses, allowing decoupling the imaging field-of-view (FOV) and imaging resolution of an optical imaging system. The imaging FOV is identical to the size of the image sensor, and the imaging resolution is typically determined by the sampling capability of the image sensor (i.e., the pixel size of sensor). By combining pixel super-resolution phase retrieval algorithms [27-29], the resolution can be further improved to sub-micron level, achieving high-throughput label-free imaging with simultaneous wide field and high resolution. However, a crucial issue in lensless imaging is the requirement for diverse measurements to enhance image quality and resolution. Strategies for capturing diverse measurements, such as moving the sample or imaging sensor to multiple axial heights [15, 30] or lateral positions [31], applying multi-wavelength [29, 32, 33] or multi-angle [34-37] illuminations, and adopting coded mask modulations [9, 26, 38], often involve precise hardware control and synchronization, which increase both the system complexity and imaging acquisition time. Additionally, the large size of the acquired images and the requirement for diverse measurements result in massive amounts of raw data, making the high-through reconstruction of lensless phase retrieval algorithms slow (typically on the order of minutes). This hinders the real-time feedback and assessment of observed samples.

To address this problem, several single-shot lensless imaging schemes based on compressive sensing or multi-beam illumination have been proposed to speed up the acquisition, but they are often limited to phase-only samples or suffer from limitations in imaging FOV and resolution [24, 39-41]. On the other hand, deep learning methods are also explored [42-46]. Rivenson et al. apply an end-to-end phase recovery approach to reconstruct high-performance complex amplitude image from a single hologram acquisition, achieving both the fast acquisition and reconstruction [42]. Chen et al. propose a Fourier imager network to further enhance the generalization of learning-based reconstruction for different samples using 3 hologram acquisitions [43]. But the generalization of these deep learning methods across different sample types and imaging system parameter settings is still limited. To improve the generalization performance, Change et al. realize the large-scale phase retrieval by combining the plug-and-play optimization framework and an enhancing neural network [44]. Self-supervised network methods [45, 46] are also explored to bypass the poor generalization of deep learning, achieving the better convergence performance such as less artifacts, higher contrast, and cleaner background, than traditional non-deep-learning methods [31, 34]. However, the data acquisition manner of the self-supervised methods remains the same as traditional methods, which needs diverse measurements, and the reconstruction speed of large-scale and high-resolution imaging are even slower.

In this work, we address the challenges of high-throughput data acquisition and phase retrieval in lensless imaging and propose a fast acquisition and reconstruction scheme for large-scale lensless microscopy (FARLS-lensless in short). We employ a multi-height lensless coherent imaging model for diverse measurements and propose a simple and continuous data acquisition strategy: manually controlling an axial translation stage to perform arbitrary movement of the image sensor, while continuously capturing multiple diffraction images at different heights with the image sensor. By eliminating the need for accurate hardware control and synchronization, we achieve the data acquisition within seconds. We further propose a high-speed parallel large-scale phase retrieval method that incorporates high-precision parameter calibration, region-wise reconstruction, region-wise auto-focusing, and parallel acceleration in the multi-height lensless model, improving the efficiency and accuracy of phase retrieval. We validate the effectiveness and practicality of the proposed method on skin section, lung tumor section, and some plant samples, demonstrating imaging performance with region-wise focusing, 7.4×5.5 mm^2^ FOV, 1.55 μm half-pitch resolution, 0.5-s acquisition time, and 44-s reconstruction time. Furthermore, by combining the pixel super-resolution principle, we achieve 1.10 μm half-pitch resolution on peripheral blood smear with the same FOV at the cost of 30-s more reconstruction time.

## 2 Methods

The whole strategy of our FARLS-lensless imaging is illustrated in Fig. 1, containing both the fast image acquisition and the fast reconstruction process. We adopt the multi-height coherent lensless imaging scheme in this work, and the image sensor is mounted on a manual translation stage instead of complex and precise motorized translation stage. The fast image acquisition is realized by arbitrarily and manually moving the axial translation stage and continuously capturing the diffraction measurements using the image sensor, requiring no precise hardware control and synchronization. In the fast reconstruction process, we first separate the full-FOV images into appropriate sub-regions with certain overlaps and then jointly use CPUs and GPUs for parallel computing to accelerate both the optical calibration step and image reconstruction calculation. We finally perform the full-FOV image stitching of all the sub-region images.

**Figure 1.**
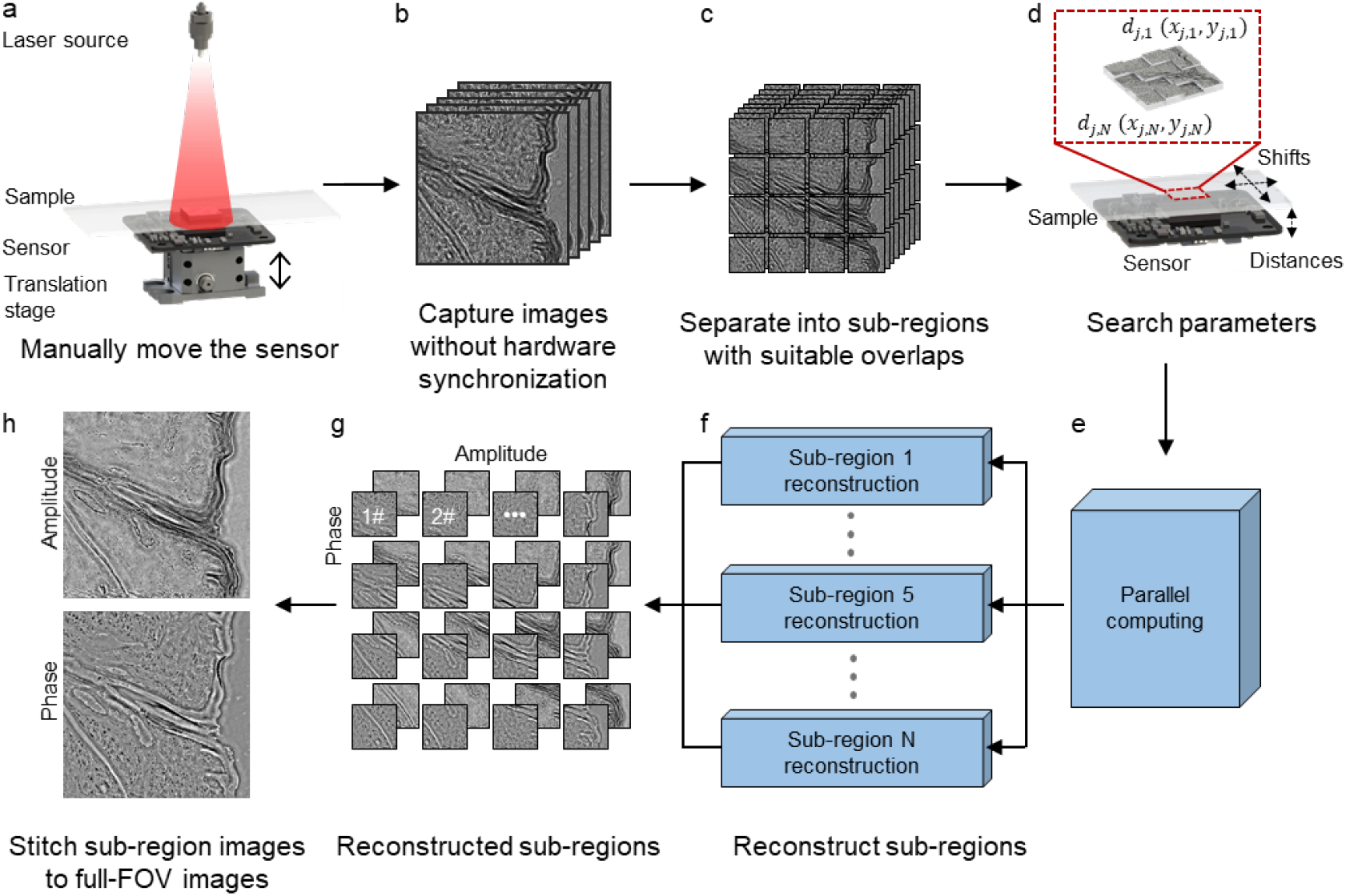
The flowchart of our proposed fast acquisition and reconstruction scheme for large-scale lensless imaging (FARLS-lensless), elaborating (a) optical set-up and image acquisition strategy, (b) multi-height raw image acquisition, (c) sub-region image separation with overlaps, (d) axial height and x-y shifts calibration, (e) parallel computing on (f) sub-region reconstruction, (g) reconstructed sub-region amplitude and phase images, and (h) full-FOV image stitching step.

### 2.1 Fast image acquisition with manual translating stage

In order to achieve fast lensless imaging, we relax the hardware accuracy requirements and realize the simple and compact imaging system, which has the potential for future portable devices. The optical setup of our system is shown in Fig. 1(a). We use the laser sources to provide the normal incidence illumination on the samples, where fiber-coupled laser sources with 632 nm wavelength (Thorlabs) and 473 nm wavelength (Changchun New Industries Optoelectronics Tech.) are respectively applied for different samples. Then the diffraction measurements of samples are captured by a monochromatic image sensor (Imaging Source DMK 33UX226, 1.85 μm pixel size and 4000×3000 pixels), which is mounted on a manual translating stage (Thorlabs).

As illustrated in Fig. 1(a-b), during the fast image acquisition step, we manually and arbitrarily perform the continuous movement of the axial translation stage (also the mounted image sensor), instead of the commonly-used precise motorized translation stage. Pre-setting a fixed exposure time, the image sensor is used to continuously capture the diffraction measurements of different axial heights, requiring no hardware synchronization. Specifically, 8 full-FOV (4000×3000 pixels, i.e., 7.4×5.5 mm^2^) raw images are continuously captured at arbitrary and unknown axial positions within 0.5 s, which is mainly limited by the sensor frame rate. Noting that the manual translating stage will inevitably introduce x-y shifts of the sample images. By utilizing high-accuracy and efficient calibration algorithms, we can calculate both the axial heights and x-y shifts with submicron-level accuracy in advance to guarantee the successful and high-quality image reconstruction. Thus, our method enables fast (0.5 s), large FOV (7.4×5.5 mm^2^), cost-effective (manual translating stage), and convenient (continuous camera shooting without hardware synchronization) lensless data acquisition for large-scale imaging.

### 2.2 Fast imaging reconstruction with region-wise parallel computing

In order to achieve high-quality and high-efficient lensless image reconstruction with the proposed compact imaging system, we realize the fast imaging reconstruction with parallel computing of FARLS-lensless imaging here. To overcome the inaccuracy control problem brought by the manually translation stage, an optimized calibration method is applied to quickly and accurately search the optical parameters, containing the axial height search and lateral shift search steps. In retrieving the information of the large-scale object based on the lensless measurements, optimized sub-region parallel calculation is employed to improve the computation speed and realize accurate region-wise focusing.

The overall reconstruction is mainly divided into three parts: image preprocessing step, optical calibration step, and phase retrieval step. The image preprocessing step separates the full-FOV captured images into sub-region images with certain overlaps, and then an optical calibration step is applied to obtain the axial positions and the lateral shifts between the object and measurements. The phase retrieval step parallelly recovers the complex amplitude information of each sub-region, and all the sub-region reconstruction results are finally stitched to obtain the full-FOV amplitude and phase images. Considering that the execution flow of the reconstruction algorithm includes both serial and parallel structures, we jointly use the CPU and GPU to further improve the computing efficiency. We make the reasonable use of CPUs to serially realize the calculation with logical sequence and utilize GPUs to simultaneously operate calculations of calibration and reconstruction among multiple sub-regions.

#### 2.2.1 Forward Imaging model

The forward model of FARLS-lensless imaging is similar to the typical coherent multi-height lensless imaging scheme. The monochromatic illumination is considered as a normally incident plane wave and can be represented as an all-one matrix *P*. The object to be reconstructed is masked as matrix *O*, and thus the light field after the object can be represented as *φ* = *P* ∙ *O*, where the ∙ is the point-wise multiplication. Then the light field is propagated from the object plane to the sensor plane, which can be calculated as the two-dimensional (2D) convolution of the point spread function (PSF) of free-space propagation and the light field after the object

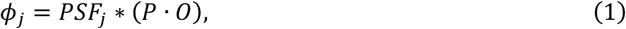

where *PSF*_*j*_ represents the PSF during the j-th image acquisition and ∗ is the 2D convolution operator. Specifically, *PSF*_*j*_ is defined as 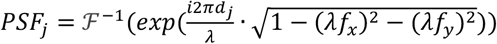, where ℱ^−1^ represents the inverse Fourier transform (IFT), *f*_*x*_ and *f*_*y*_ respectively represent the spatial frequency along x and y directions, *λ* represents the illumination wavelength, and *d*_*j*_ denotes the object-to-sensor distance in the j-th acquisition. In real calculation, the 2D convolution operation is performed in Fourier plane for high efficiency. Finally, the j-th captured raw image *I*_*j*_ can be represented as

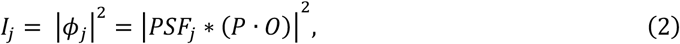

where | ∙ | is the modulus operator. In total, 8 raw images *I*_*j*_ (*j* = 1: 8) are captured during the image acquisition process of a certain sample.

#### 2.2.2 Optical calibration step

To eliminate the influence on reconstruction caused by the unknown and random motions of the image sensor, we need to perform a calibration step to determine the x-y-z shift parameters of the object relative to the sensor acquisition. We separate each captured full-FOV raw image into N sub-region images in advance for parallel computing to improve the computational efficiency, as illustrated in Fig. 1(c). Adjacent sub-region images have some overlaps, which are about 10% of the sub-region image size, to guarantee the successful full-FOV stitching later. Then we optimize and apply a calibration algorithm to search the optical parameters of all these N×8 images. As mentioned above and shown in Fig. 1(d), the calibration step contains searching the axial heights (i.e., the distance between the sensor and the corresponding sub-region object) and the lateral shifts (i.e., the x-y shifts between the sub-region image relative to the corresponding sub-region object) of N×8 images. Noting that a large FOV sample generally has different in-focus positions in different regions. Benefiting from the sub-region calibration, we can obtain the accurate region-wise focusing consequently.

##### Axial height search

We denote the n-th sub-region image (*n* = 1: *N*) in the j-th captured raw image (*j* = 1: 8) as *I*_*j,n*_ and the corresponding height as *d*_*j,n*_. Although the full-FOV sample has different in-focus positions in different regions, we consider that this difference is global among 8 raw acquisitions, which means all sub-regions share the same height differences ∆_*j*_= *d*_*j,n*_ − *d*_1,*n*_(*j* = 1: 8). Thus, we first choose a reference sub-region and search the distances *d*_*j,ref*_ of all 8 images *I*_*j,ref*_ inside the reference sub-region. After that, the height parameters of other sub-regions can be determined by searching for the elevation differences between their first images and the corresponding first image in the reference sub-region. By this, we can largely shorten the height searching time.

In searching all the heights *d*_*j,ref*_ (*j* = 1: 8) of the reference sub-region, we firstly parallelly search the minimum axial height *d*_1_,_*ref*_ and the maximum axial height *d*_8_,_*ref*_. Then we use *d*_1_,_*ref*_ and *d*_8_,_*ref*_ to further narrow down the search ranges for other 6 heights. The height searching algorithm [47] in use utilizes the concept of exhaustion and the “sparsity of the gradient” (SoG) to find out the optimal height from a given rough range. Taking the determination of the minimum axial height *d*_1 *ref*_ as an example, we denote the initial range of *d*_1 *ref*_ as a vector consisting of 2*M* +1 values as 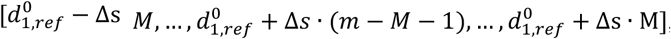, where *m* represents the m-th given height in the roughly estimated range, and 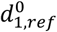 represents the central value of the range. ∆s denotes the search step, which we typically set as 100 μm in the beginning. The complex amplitude 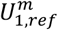 is obtained by propagating *I*_1_,_*ref*_ back at distance 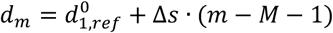, and the corresponding SoG is defined as

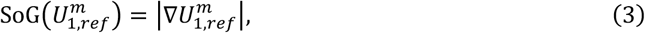

where ∇(∙) is the gradient operator. 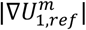 represents applying the modulus operation on the gradient of 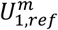, which can be calculated as 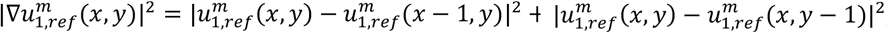, where 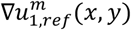 and 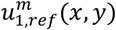 represent the values in the coordinate (*x, y*) of 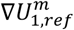 and 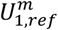 respectively. Then “Tamura of the gradient” (TC) of the given height *d*_*m*_ is defined as

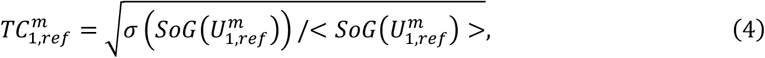

where σ(∙) is the standard deviation and <∙> is the mean operator. We calculate all the 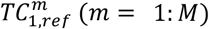 and regard the 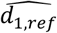 corresponding to the maximum *TC* value as the accurate estimation of the targeted *d*_1_,_*ref*_. Then we denote this 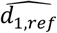 as the new central value 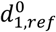, and reduce the search step ∆ (usually is reduced by one order of magnitude) to start a new round of search. We repeat the above process until the required accuracy of *d*_1_,_*ref*_ is achieved, and we select 1 μm degree of accuracy in this work. We use the same process to determine the value of *d*_8_,_*ref*_, which is calculated parallelly with *d*_1_,_*ref*_.

After *d*_1_,_*ref*_ and *d*_8_,_*ref*_ are both determined, we narrow down the search range for other 6 heights in this reference sub-region for quicker search. We begin with a rough assumption that the height interval of each image is almost equal during the sensor acquisition process, so the initial guesses of central values 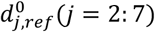 are obtained by uniformly sampling the range [*d*_1*ref*_, *d*_8_,_*ref*_]. Then the same height searching algorithm is applied in parallel to search the targeted heights *d*_*j*_,_*ref*_ (*j* = 2: 7) within the range 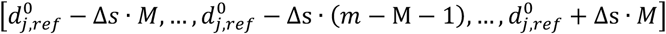, where the step ∆*s* can be initialized as 10 μm now, which significantly narrows the search range and shortens the computing time. We repeat the above process until the 1 μm accuracy of *d*_*j*_,_*ref*_ (*j* = 2: 7) is achieved. So far, all heights *d*_*j*_,_*ref*_ (*j* = 1: 8) of the reference sub-region are determined. ∆_*j*_ can be calculated accordingly as ∆_*j*_= *d*_*j*_,_*ref*_ − *d*_1_,_*ref*_ (*j* = 1: 8) in the reference sub-region and considered global unchanged as above mentioned.

Subsequently, we search the height differences ∆*d*_1,*k*≠*ref*_ between the first image *I*_1_,_*ref*_ in the reference sub-region and the first images in all other sub-regions *I*_1,*k*≠*ref*_ (*k* = 1, …, *ref* − 1, *ref* +1, …, *N*). Using the already known height *d*_1_,_*ref*_ as the initial central height, the height searching algorithm is again employed in parallel to fast calculate the height differences ∆*d*_1,*k*≠*ref*_ between the *I*_1,*k*≠*ref*_ and *I*_1_,_*ref*_. Then the heights of first images in all other sub-regions can be calculated as *d*_1,*k*≠*ref*_=Δ*d*_1,*k*≠*ref*_ +*d*_1_,_*ref*_. Finally, we can directly obtain the axial heights of all other sub-region images by *d*_*j,k*≠*ref*_ = *d*_1,*k*≠*ref*_+∆_*j*_= ∆*d*_1,*k*≠*ref*_ +*d*_1_,_*ref*_ +*d*_*j, ref*_ − *d*_1_,_*ref*_ = ∆*d*_1,*k*≠*ref*_ +*d*_*j*_,_*ref*_ (*j*= 1: 8, *k* = 1, …, *ref* − 1, *ref* +1, …, *N*). By this, the axial heights of all sub-regions are accurately calibrated. The detailed processes of axial height search are illustrated in Fig. S1.

##### Lateral shift search

Given the certain sub-region images and their axial height parameters, we can parallelly calculate the lateral shifts of each sub-region images (with respect to the first image *I*_1,*n*_) using a registration algorithm based on Fourier cross correlation [48]. We first back propagate a certain captured sub-region image *I*_*j,n*_(*j* = 2: 8) and the first image *I*_1,*n*_ (as the reference image) to their object plane, obtaining *U*_*j,n*_ and *U*_*j*,1_ at distance *d*_*j,n*_ and *d*_*j*,1_ respectively. We then calculate the Fourier transform (FT) of |*U*_*j,n*_ |and |*U*_1,*n*_ | to find the peak location of the cross correlation of these two images, namely the targeted shifts. The cross-correlation *R*_*j,n*_ of the reference image and the searched image can be obtained by

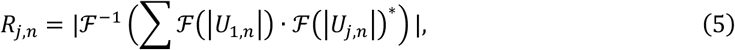

where ℱ(∙) represents FT, and (∙)^∗^ represents the complex conjugation. The coordinate corresponding to the maximum value of *R*_*j,n*_ is the targeted shift (*x*_*j,n*_, *y*_*j,n*_). We can use the similar steps to calculate the lateral shifts of all sub-region images with multiple heights.

#### 2.2.3 Parallel phase retrieval

Based on the sub-region raw images and calibrated parameters, we realize the parallel computation for fast and large-scale lensless phase retrieval based on the extended ptychographic iterative engine (ePIE) [31]. A certain number of parallel workers are created to perform the reconstruction, where each sub-region raw images are input into an individual worker. The number of workers is set according to the core numbers of CPU, fully utilizing the computing resources. The phase retrieval of the full-FOV image are separated into the N independent parallel tasks (i.e., N sub-region images).

In each task, several iterations (*l* = 1:*iter*) are performed to iteratively update the object information inside a sub-region. Each iteration contains several sub-iterations (*j* = 1: 8), which corresponds to update the object using 8 raw images with different heights. We input the illumination *P* and the initial guess of the object 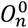. In each sub-iteration, we obtain the light field on the sample plane by *φ*_*j,n*_ = *P* ∙ *O*_*j,n*_. Based on the height and shift parameters, we can predict the light field *ϕ*_*j,n*_ following the Eq. (1) and the captured image 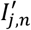 on the sensor plane following the Eq. (2). We then use the real-captured corresponding image *I*_*j,n*_ to update the light field on the sensor plane and obtain the updated light field 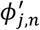. We next back propagate the 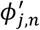 to the object plane and get the corresponding light field 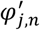. We finally update the object distribution 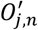 based on 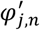^′^ and previously estimated object distribution 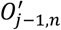. The update steps can be written as:

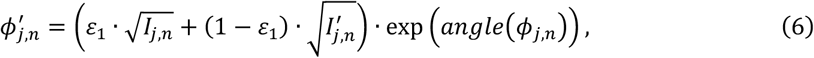

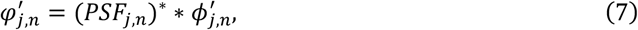

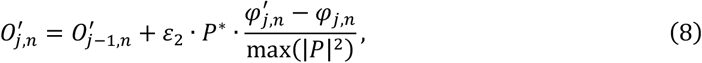

where *angle*(∙) returns the phase angle, and *ε*_1_ and *ε*_2_ are preset parameters. We traverse all heights and successively update the object using multi-height images using the above steps to finish a whole iteration. After doing several iterations, the object information of a certain sub-region is recovered. The phase retrieval of each sub-region is parallelly executed in GPU, which has a high calculation efficiency. We finally stitch all sub-region recovered images to a full-FOV image of the object. Our parallel image reconstruction process is operated in MATLAB using a desktop computer with Intel(R) Core(TM) i9-9900X CPU @ 3.50GHz, 128GB of RAM, and GeForce RTX 2080 Ti (Nvidia).

We summarize the whole reconstruction process of our strategy in the Fig. 2.

**Figure 2.**
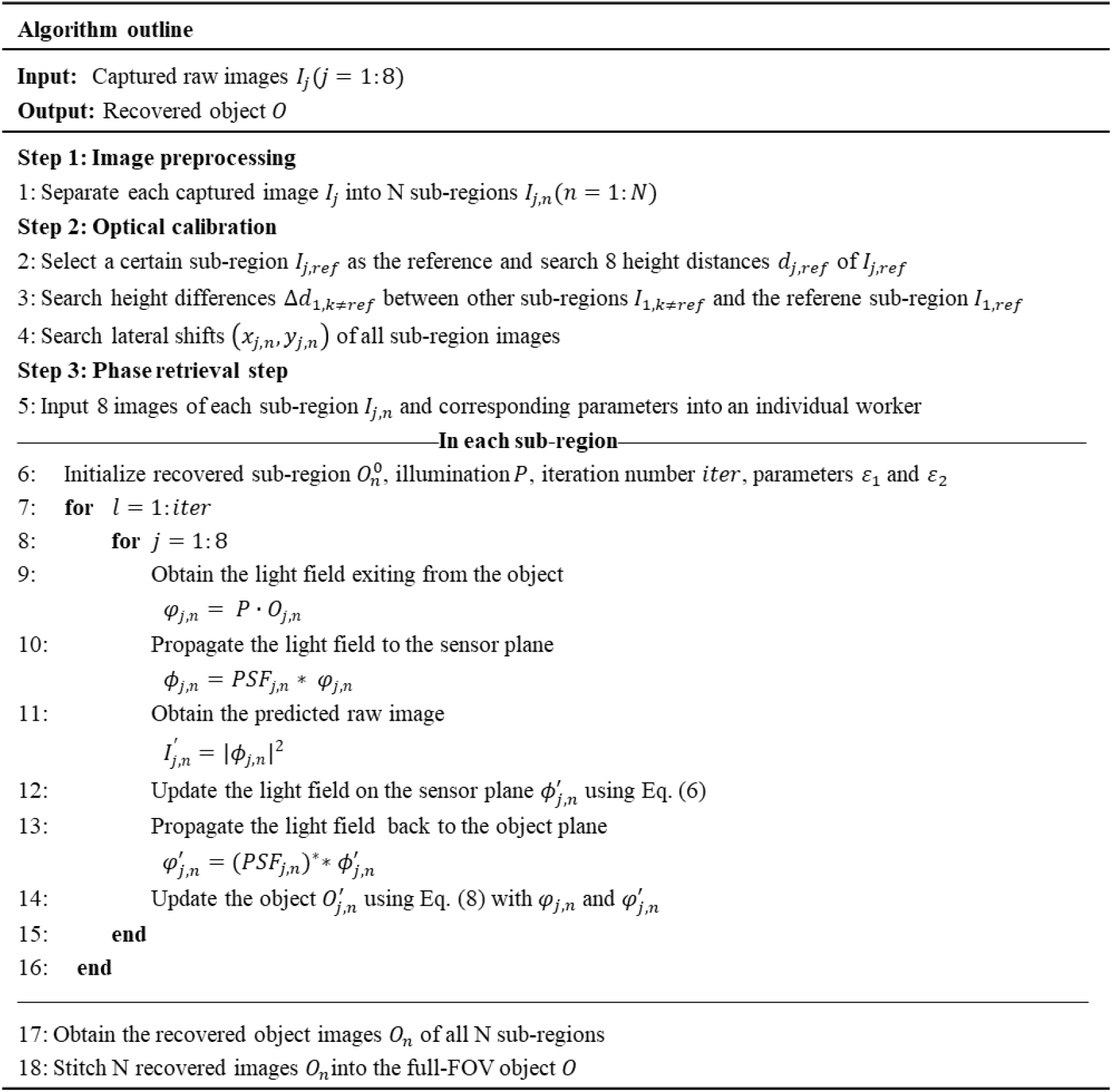
Algorithm outline of fast imaging reconstruction with parallel computing.

## 3 Results

### 3.1 Reconstruction accuracy validation by arbitrary and manual height movement

We first validate the effectiveness and feasibility of our proposed fast image acquisition strategy, i.e., the manual height modulation of the sample diffraction information. For comparison, we build up a conventional multi-height lensless imaging system using a fiber-coupled laser source with 632 nm wavelength, the same monochromatic sensor with 1.85 μm pixel size, and a motorized translation stage (Thorlabs). As shown in Fig. 3, we respectively capture 8 raw images (2400×2400 pixels) of the fig fruit sample (commercially bought from AOXING Laboratory Equipment) using both acquisition strategies, search required parameters by the calibration step, and reconstruct complex amplitude (both amplitude and phase) images using the proposed algorithm. The conventional strategy needs to guarantee the slow and precise motorized motion of the image sensor (Fig. 3(a)), where we move the sensor with 50 μm step each time. On the contrary, our strategy only performs the quick and unprecise motion of the image sensor by using a manual translation stage (Fig. 3(e)). After reconstruction, two different strategies obtain similar performance in both the amplitude and phase imaging, as shown in Fig. 3(b) and 3(f) respectively. Some close-up images are also exhibited to better compare the reconstruction results, where we can find that the fine structures of the sample are well resolved in both situations. It demonstrates that our fast acquisition strategy will not degrade the reconstruction quality. Preserving the high imaging quality, our strategy not only reduces the system complexity and cost, but also shorten the acquisition time from tens of seconds to approximate 0.5 s. It is because the conventional multi-height image acquisition needs to a precisely motorized translation stage to move the sensor to a specific position each time, and waits for the image sensor to capture the certain raw image, i.e., it needs the hardware synchronization between the sensor and the translation stage. But our strategy gets rid of the motorized translation stage and the hardware synchronization, benefiting from the proposed calibration and reconstruction algorithms.

**Figure 3.**
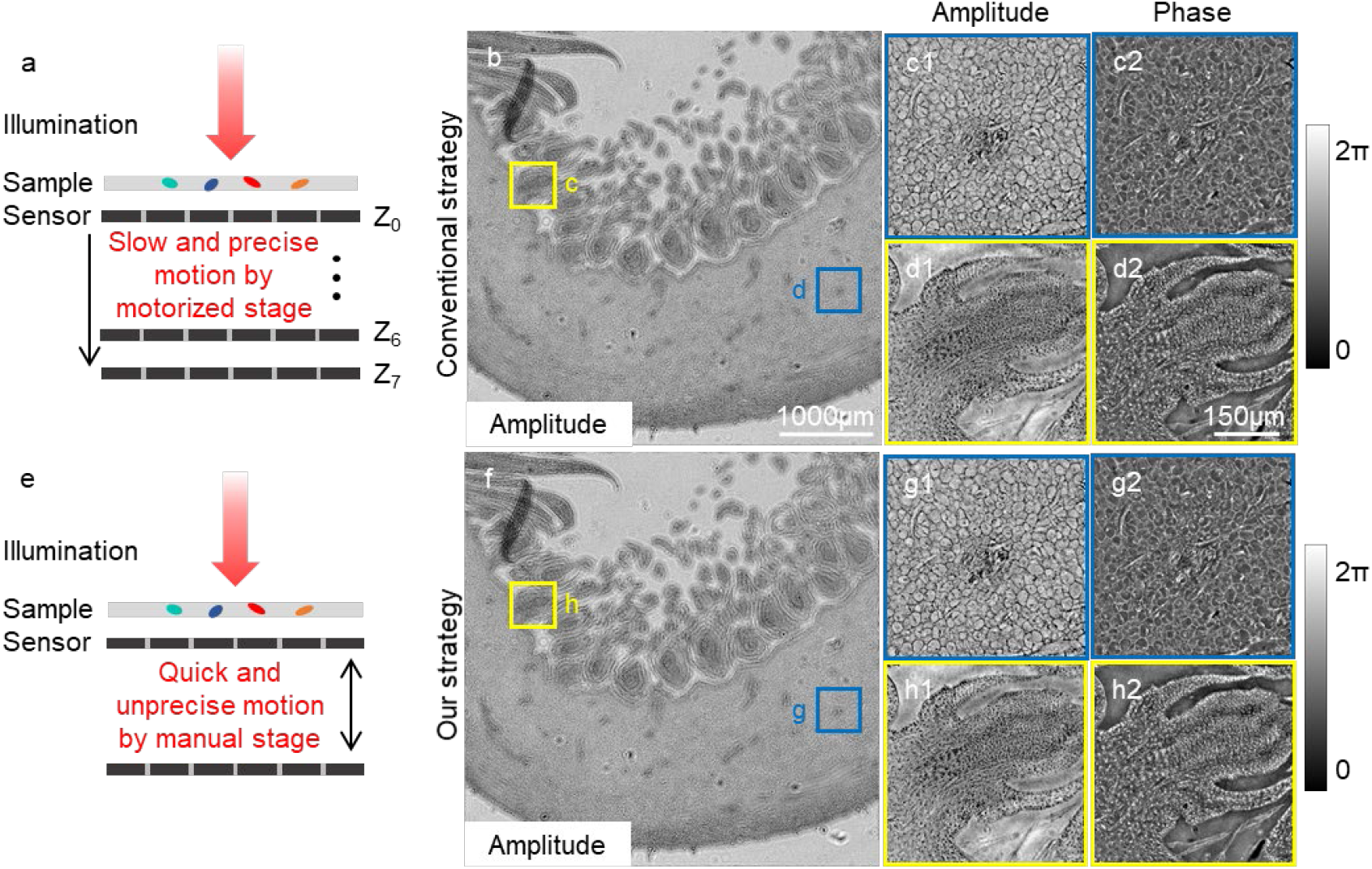
We validate the feasibility of the proposed fast image acquisition on a fig fruit sample. (a) Conventional multi-height image acquisition mounts the image sensor on a motorized stage, and precisely move the sensor to multiple axial positions to capture 8 raw measurements (2400×2400 pixels each), taking tens of seconds acquisition time. (b) The corresponding reconstructed amplitude image using the conventional image acquisition and (c-d) some close-up amplitude (c1, d1) and phase images (c2, d2) highlighted in (b). (e) Our fast image acquisition uses a manual translation stage to replace the motorized translation stage, and arbitrarily moving the image sensor. We then continuously capture the 8 raw images for imaging recovery within 0.5 s. (f) The corresponding reconstructed amplitude image using the proposed image acquisition and (g-h) some close-up amplitude (g1, h1) and phase images (g2, h2) corresponding to the highlighted areas in (f). From the results, we find that these two different strategies obtain similar performance in both the amplitude and phase imaging. Fine structures are well resolved in both situations, demonstrating that our fast acquisition preserves the high imaging quality and shortens the acquisition time to less than one second.

We also test our acquisition strategy by imaging the USAF-1951 resolution chart and the reconstructed images show that our strategy can achieve the same resolution as the conventional strategy (Fig. S2), demonstrating that our strategy can acquire and resolve the same sample information as the conventional strategy.

### 3.2 Resolution evaluation

In this section, we test the achieving resolution of the proposed method by imaging the USAF-1951 resolution chart and microspheres (3-μm dimeter, polystyrene microsphere). We can also integrate the pixel super-resolution (PSR) algorithm [28] into the original reconstruction to achieve a higher resolution, scarifying a little more computation time. To perform the pixel super-resolution algorithm, we modify the update step (step 12 in the algorithm outline) of the original reconstruction algorithm in the sub-sampled way [28]. Specifically, the updated light field on the sensor plane is calculated as 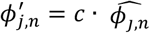, where the update weight *c* is calculated as 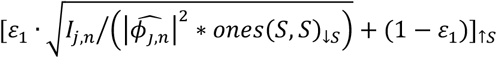, and ↑ *S* and ↓ *S* represent *S* by *S* up-sampling and down-sampling.

During both PSR and no PSR reconstructions, we input an all-one matrix as the initial guess of the object and perform the same number of iterations. We also capture 8 raw images by our proposed fast acquisition strategy and recover the complex amplitude information of samples. The no PSR algorithm can resolve 1.55 μm line width in the USAF-1951 resolution chart (group 8 element 3), which is little higher than the sampling limit of the sensor pixel size (1.85 μm), as the results shown in Fig. 4(a). The PSR algorithm can achieve higher resolution, resolving 1.10 μm line width in the USAF-1951 resolution chart (group 8 element 6), as the results shown in Fig. 4(b). The achieving resolution is also demonstrated by imaging 3-μm microspheres, where Fig. 4(c) and 4(e) compare the recovered amplitude images of two algorithms, and Fig. 4(d) and 4(f) exhibit the normalized line profiles along the red line in corresponding images. PSR algorithm shows an obvious resolution improvement based on the Rayleigh Criterion, where adjacent microspheres can be solved more clearly. Noting that the PSR algorithm shares the same calibration step with the non-super-resolution one, but it consumes a little longer reconstruction time of phase retrieval, which is related to the size of raw images. We will discuss the time consumption in the following Section 3.3.

**Figure 4.**
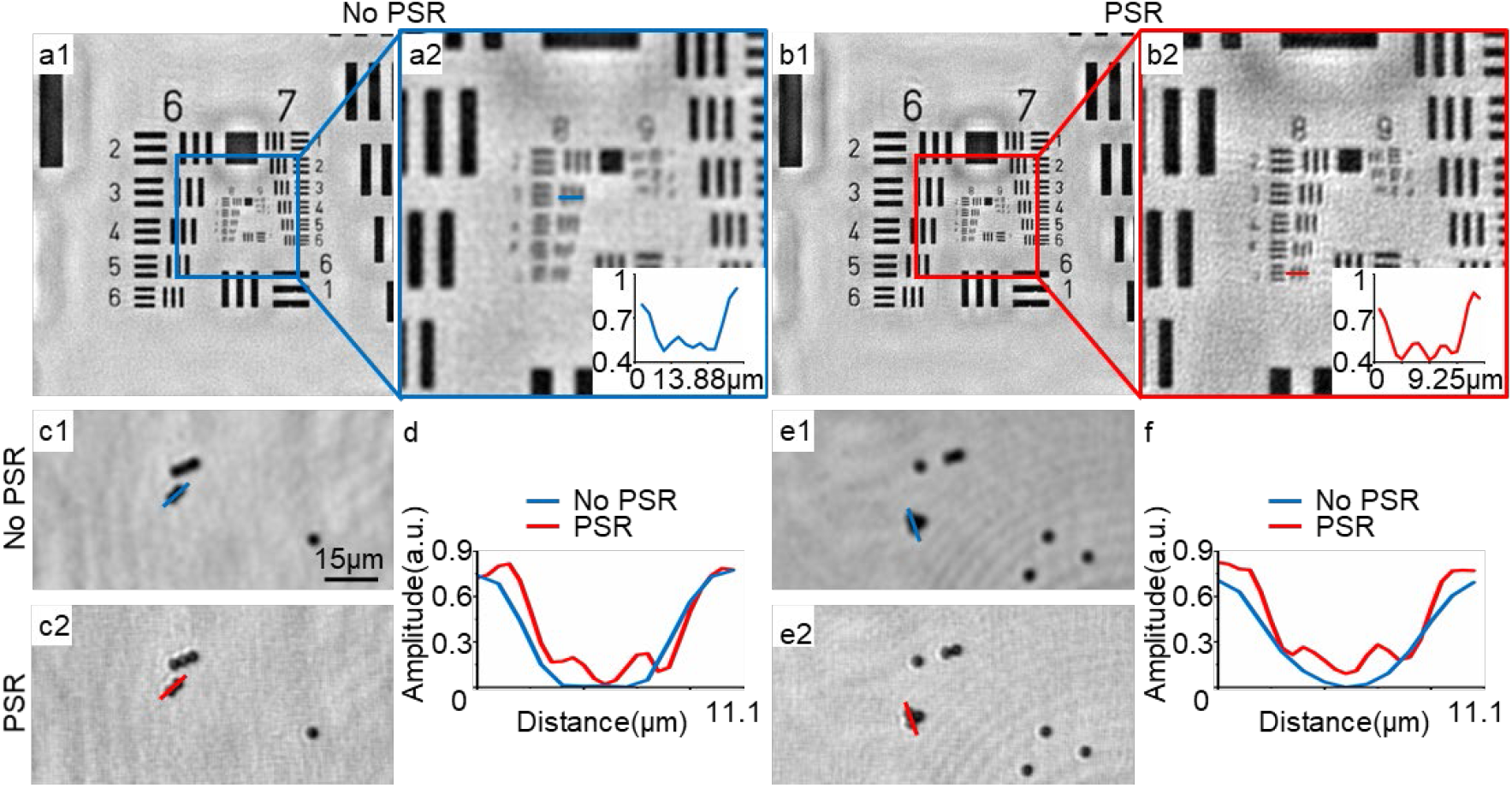
We test the achieving resolution of our method by imaging the USAF-1951 resolution chart and microspheres. (a1) Reconstructed resolution chart image using the proposed algorithm with no pixel super-resolution (no PSR) and (a2) its close-up image. (b1) Reconstructed resolution chart image using the pixel super-resolution (PSR) algorithm and (b2) its close-up image. Through visual observation and the normalized line profile in (a2), the no PSR algorithm can resolve group 8 element 3 of the resolution chart, achieving 1.55 μm half-pitch resolution. Through the results in (b2), the PSR algorithm can resolve group 8 element 6 of the resolution chart, achieving 1.10 μm half-pitch resolution. We also demonstrate the resolution improvement of PSR method by imaging 3-μm diameter microspheres, where the no PSR reconstructed microsphere images are shown in (c1) and (e1) and the PSR reconstructed microsphere images are shown in (c2) and (e2). We can witness significant resolution improvement by the recovered images and the normalized line profiles shown in (d) and (f) respectively.

### 3.3 Full-FOV imaging with parallel region-wise focusing

Lensless imaging inherently enjoys the advantage of large FOV, which is equivalent to the size of image sensor. However, the biological samples typically have varying focus planes among, due to the thickness or the uneven placement of samples different imaging area. Here in this section, we demonstrate that, besides the fast computation speed, our parallel sub-region calibration and reconstruction naturally possess the region-wise focusing ability, which is very beneficial for full-FOV imaging.

We first use a 632 nm laser to illuminate and reconstruct the full-FOV human skin sample (commercially bought from KONUS CORP.), as shown in Fig. 5. The reconstructed full-FOV amplitude and phase images are respectively shown in Fig. 5(a) and 5(e), which have about 7.4×5.5 mm^2^ FOV. We compare the results with our region-wise focusing and those with globally-consistent focusing in Fig. 5(b-d) and 5(f-h), which are the magnified images of the highlighted areas in the full-FOV images. Specifically, the results of the globally-consistent focusing are shown in Fig. 5(b1-d1) and 5(f1-h1), while the results recovered using our region-wise focusing are shown in Fig. 5(b2-d2) and 5(f2-h2), with the recovered focusing positions given in the top right corner. From the results, our method resolves much sharper and clearer structures in both the amplitude and phase images indicated by the boxes and arrows, overcoming the defocus artifacts. We also calculate and show the fitted elevation map of the full-FOV skin sample in Fig. S3 to better present the focusing positions of each sub-region. Noting that the data acquisition time of 8 full-FOV raw images (each has 4000×3000 pixels) is within 0.5 s. Besides, our full-FOV reconstruction with parallel region-wise focusing consumes about 44 s, containing 2-s image preprocessing step, 21-s calibration step, 20-s reconstruction step, and 1-s stitching step.

**Figure 5.**
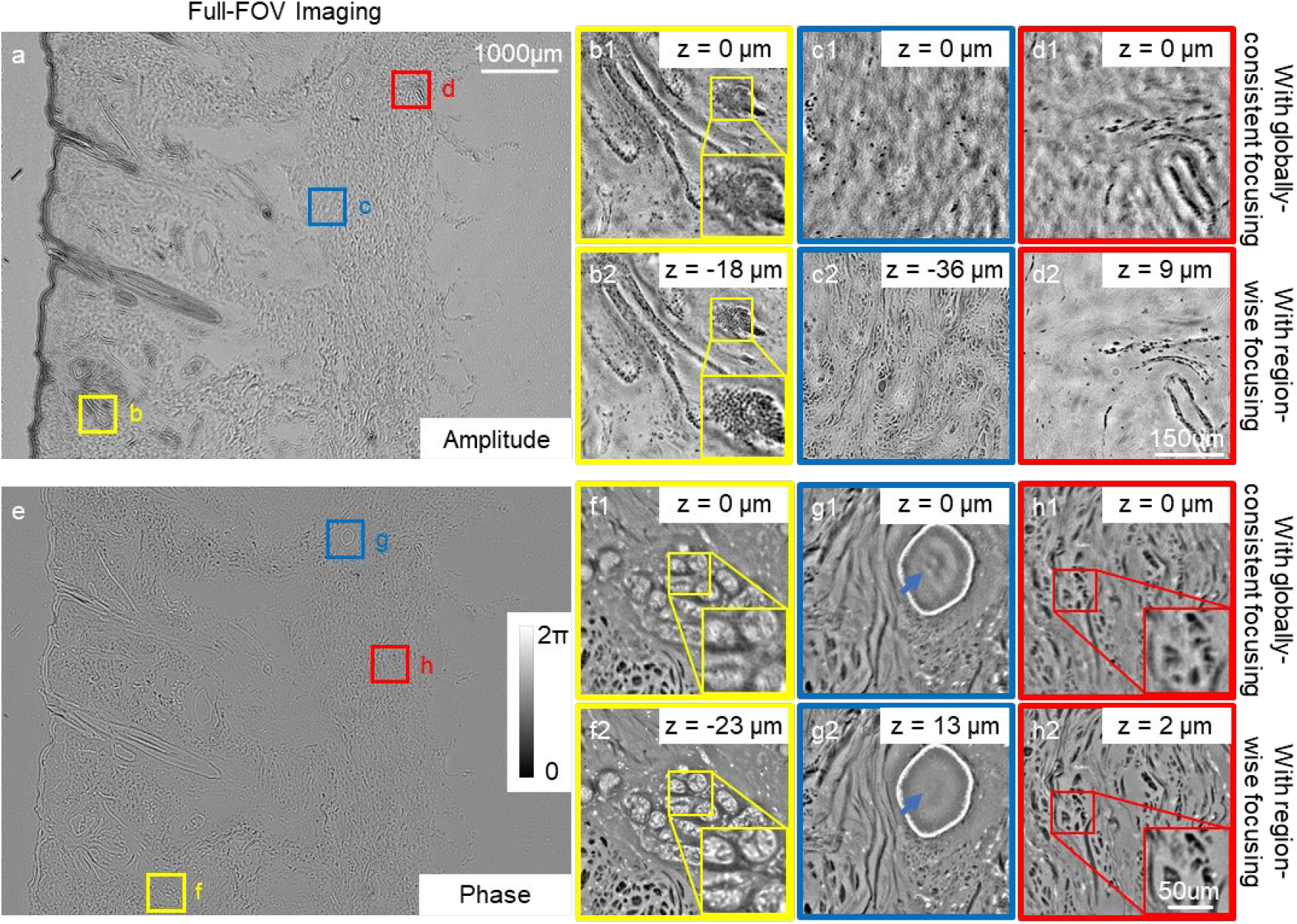
We exhibit the large-scale imaging ability with the region-wise focusing of our method on the human skin sample. The stitched full-FOV (7.4×5.5 mm^2^) amplitude (a) and phase (e) images. We further show some close-up images marked by color boxes in (a) and (e) to better show the region-wise focusing ability of our method, as presented in (b-d) and (f-h). Without region-wise focusing (i.e., with globally-consistent focusing), the recovered images are full of defocus blurs, as show in (b1-d1) and (f1-h1). In comparison, our reconstruction with region-wise focusing resolves much sharper and clearer structures in both the amplitude and phase images, as presented in (b2-d2) and (f2-h2). The recovered focusing positions are given in top right corner of close-up images. Arrows and boxes further highlight the image quality improvement by our method. Our method needs less-than-0.5-s data acquisition time and 44-s parallel reconstruction time.

We then use a 473 nm laser to illuminate and image the lung tumor section to further verify that our method has the large FOV imaging and region-wise focusing ability. We respectively exhibit the full-FOV reconstruction results with region-wise focusing, the fitted elevation map, and the comparison with 5× bright-field microscopy of the sample in Figs. S4-S6. We also test the digital staining performance [49] on the recovered amplitude image of the tumor section in Fig. S6, which can be utilized for further quantitative analysis.

### 3.4 Full-FOV pixel super-resolution imaging of blood smear

In this section, we perform the PSR algorithm on the label-free peripheral blood smear for higher resolution full-FOV imaging by consuming more computation time. We still acquire 8 full-FOV raw images with manually multi-height modulation under the 473 nm laser illumination, taking approximate 0.5-s acquisition time. We apply the parallel PSR reconstruction algorithm to quickly recover the image of blood cells, as the full-FOV amplitude and phase images shown in Fig. 6(a-b). For better comparison, we respectively show some close-up images highlighted in Fig. 6(a-b), which are recovered in different conditions, including the region-wise focusing with no PSR in Fig. 6(c1-h1), globally-consistent focusing with PSR in Fig. 6(c2-h2), and region-wise focusing with PSR in Fig. 6(c3-h3).

**Figure 6.**
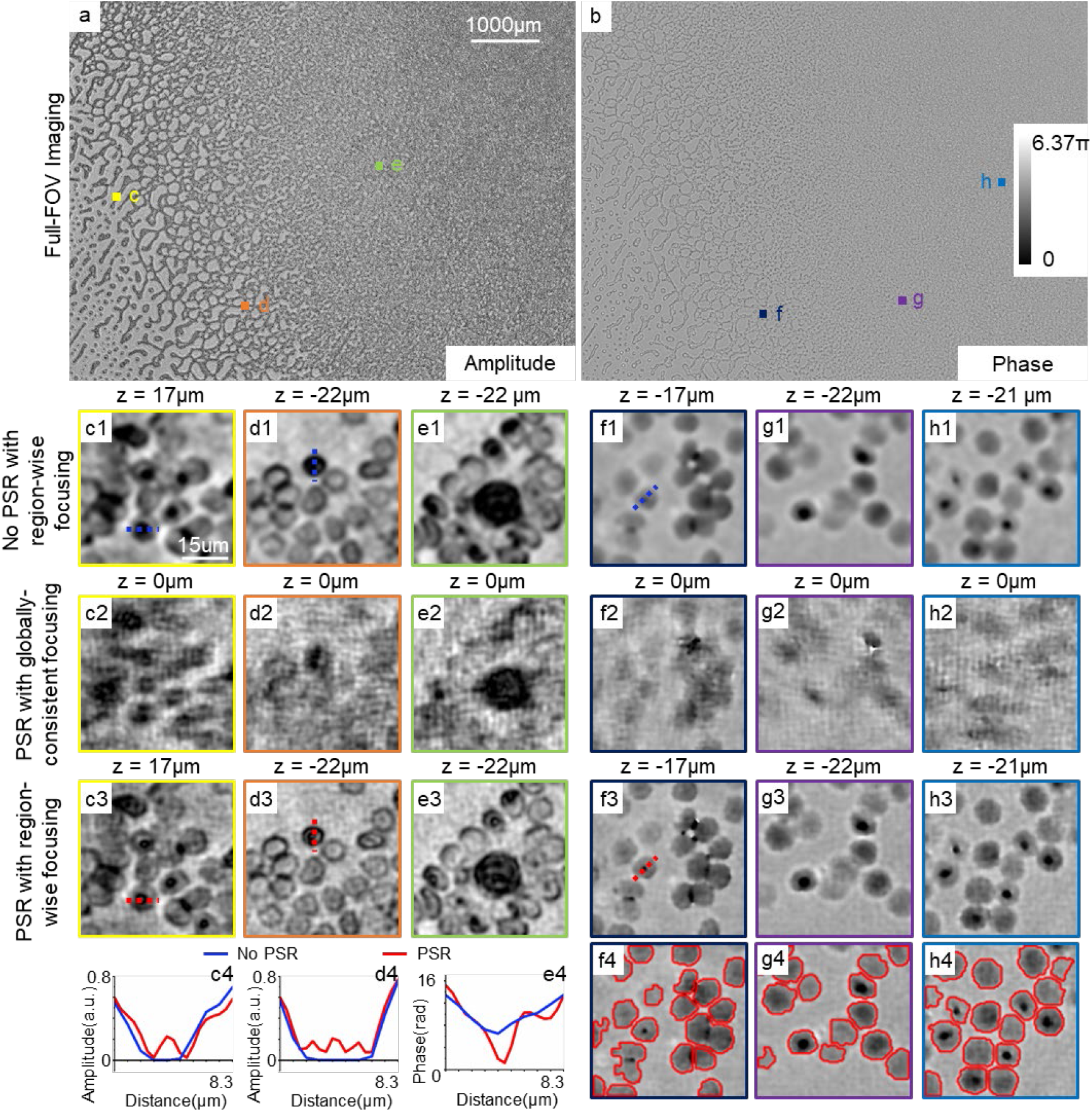
We demonstrate the pixel super-resolution (PSR) reconstruction of full-FOV imaging of the peripheral blood smear. (a) Reconstructed full-FOV (7.4×5.5 mm^2^) amplitude and (b) phase images. By comparing with the results of no pixel super-resolution (no PSR) with region-wise focusing (c1-h1) and the results of PSR without region-wise focusing (c2-h2), we better exhibit the imaging performance of PSR with region-wise focusing (c3-h3). From these magnified imaging results, which are highlighted regions in (a) and (b), the biconcave-disk-like shapes of red blood cells can well be resolved by PSR with region-wise focusing only, demonstrating its resolution improvement ability. This resolution improvement can also be witnessed in the normalized line profiles in (c4-e4). The no PSR with region-wise focusing method takes about 44-s computation time for the full-FOV imaging., while the PSR with region-wise focusing method takes about 76 s computation time, obtaining approximate 1.41 times resolution improvement. We also conduct the cell segmentation of blood cells based on the high-quality phase imaging (f4-h4).

There is no doubt that the globally-consistent focusing with PSR in Fig. 6(c2-h2) has the worst performance (showing heavily defocus blurs), since the focus depths are inaccurately estimated in this condition. The region-wise focusing with no PSR in Fig. 6(a-b) uses the same reconstruction process as the full-FOV imaging results in Section 3.3, taking about 44-s computation time. Although with the region-wise focusing ability, the biconcave-disk-like shapes of red blood cells can’t be resolved clearly due to the limited resolution. In comparison, the region-wise focusing with PSR in Fig. 6(c3-h3) can well obtain the fine details of blood cells, successfully resolving the biconcave-disk-like shapes. This resolution improvement can also be witnessed in the normalized line profiles between no PSR and PSR in Fig. 6(c4-e4), which are marked by red dash lines in Fig. 6(c3), 6(d3), and 6(f3).

The PSR reconstruction process approximately improves 1.41 times resolution (testing by the resolution chart imaging in Fig. 4 above) by consuming about 30-s more computation time, which totally takes 76 s reconstruction time for the full-FOV imaging. We further conduct the cell segmentation of blood cells based on the high-quality and high-resolution phase imaging reconstructed by the PSR algorithm with region-wise focusing, as shown in Fig. 6(f4-h4) and Fig. S7. Specifically, we apply a classical three-step segmentation method [50] here, containing the binarization, morphology processing, and watershed segmentation steps. The segmentation results of oral epithelial cells are also presented in Fig. S8, as an additional example. Noting that we need to make a compromise between resolution and reconstruction time in our method by choosing no PSR or PSR, and we recommend selecting a more suitable resolution for imaging different samples.

## 4 Discussion and Conclusion

In this work, we propose the FAFRLS-lensless method to address the low-efficient acquisition and reconstruction problems in large-scale lensless imaging. We apply the accurate position parameter calibration to relax the requirement for high-precise control and modulation of raw image acquisitions. We thus can adopt a manual and fast multi-height modulation of sample diffraction measurements for image acquisition within seconds. For high-quality and full-FOV lensless recovery, we implement both the efficient sub-region calibration and reconstruction by utilizing the parallel computation structure, taking only about 44-scecond time for 7.4×5.5 mm^2^ FOV imaging. We further integrate the PSR algorithm into our method to obtain 1.41-time higher resolution at the cost of 30-s more computation time for full-FOV imaging. Through real experiments of resolution chart and microspheres, we demonstrate that our method can achieve high-quality lensless imaging using both no PSR and PSR algorithm, achieving 1.55 μm and 1.10 μm half-pitch resolutions respectively. We also prove that the proposed method can be used to conveniently and effectively image the plant samples, human skin samples, lung tumor sections, and blood smears, possessing the region-wise focusing ability and resolving the fine structures of samples. Based on high-accuracy reconstruction, we realize exemplary cell segmentation and digital staining experiments. Our method inherits the large-scale imaging ability of lensless imaging, and it is simultaneously equipped with the high acquisition and reconstruction speed. It only requires the low-priced manual translation stage with no hardware synchronization needed. Therefore, it is well-suited for low-cost and high-speed scanning of biomedical smears and sections, benefiting for the following high-throughput screening and pathological research. Future researches rely on further improving the performance of cell segmentation and digital staining, as well as further increasing the imaging resolution or accelerating the reconstruction speed through the hardware and algorithm improvement.

## Acknowledgments

The work was supported by the National Science Foundation of China (NSFC) (Grant Nos. 62071219 and 62025108). X. Cao and Y. Zhou supervised this project. Y. Zhou and B. Xiong conceived and designed the experiments. W. Song, L. Wu, and L. Fan prepared the samples. W. Song conducted the experiments. Y. Zhou, W. Song, and J. Wang realized the algorithms. W. Song, B. Xiong, and S. Jiang analyzed and interpreted the experimental data. All authors discussed the results and contributed to the final manuscript.

## Data Availability Statement

The data and codes that support the figures and findings within this article are available from the corresponding authors upon reasonable request.

## Appendix A

### Sample Preparation

#### Polystyrene microspheres

Polystyrene microspheres of 3-μm diameter (M122075, aladdin) was diluted 2500 times with distilled water and smeared on a 1-mm-thick glass slide. The microspheres were imaged at room temperature.

#### Peripheral blood smears

The EDTA-2K anticoagulant whole blood samples were obtained from the clinical laboratory at Zhongda Hospital Southeast University. Aliquots of blood samples were processed into blood smears and stained with Wright-Giemsa, utilizing a Mindray SC-120 Automatic Slide Maker (Shenzhen Mindray Corporation, China).

#### Lung tumor section

The lung tumor sections were heated at 90°C, then dewaxed with dimethylbenzene and ethanol of gradient concentration (95%, 90%, 80%, 70%) for 10 min each, and last processed with 3% H_2_O_2_ and citrate retrieval solution (Guoyao chemical reagent co., LTD). The sections were incubated with primary antibodies after the above procedure at 4°C overnight and washed with PBS 3 times. Then the sections were labeled by the HRP labeled secondary antibodies at 37°C for 30 min and stained with DAB for 2 min. After being washed with PBS, the sections were dehydrated with ethanol of gradient concentration (70%, 80%, 90%, 95%).

#### Oral epithelial cells

Oral epithelial cells, taken from volunteers, were diluted with distilled water and smeared on a 1-mm-thick glass slide. We imaged oral epithelial cells sample at the room temperature.

